# VPF-Class 2.0: a taxonomy-centered framework for automatic viral classification

**DOI:** 10.64898/2026.03.20.713201

**Authors:** Luis J. Vidal, Joan Carles Pons, Mateus B. Fiamenghi, Nikos Kyrpides, Mercè Llabrés

## Abstract

Rapid expansion of viral sequence data demands classifiers that scale, track ICTV updates, and provide interpretable evidence. We present VPF-Class 2.0, an updated successor to VPF-Class, centred on the taxonomic classification, that retains marker-driven protein domain detection but replaces rule-based voting with a lightweight supervised model on per-genome marker-composition features. In controlled benchmarks, VPF-Class 2.0 achieves near-perfect family-level performance and strong genus-level accuracy while increasing confident annotation coverage. Under a practical confidence threshold (0.3), performance improves and matches or exceeds representative tools within shared taxonomic scopes. We further introduce an interpretability study that relates errors to the genus specificity of activated markers. Finally, we demonstrate applicability on large real-world viromes with consistent labels and substantial agreement with graph-based classifications. The implementation of VPF-Class 2.0 can be downloaded from https://github.com/luisvidalj/VPFClass2.git.

## Introduction

Viruses are the most abundant biological entities on Earth, infecting hosts across all domains of life and shaping microbial mortality, gene flow, and ecosystem processes (see [1, 9]).

Viral diversity differs from cellular life in two key ways [7]: they employ all known genome types and replication strategies in the Baltimore framework [8], and they evolve at high rates that allow exploration of vast sequence space, to the point that a single viral family can exceed the genetic diversity of entire cellular domains [10].

Among complementary organizational criteria (genome type/replication, virion morphology/host range, ecological signals), we focus on taxonomy, an evolutionary framework that enables consistent communication and comparison across the expanding virosphere.

Viral taxonomy is a dynamic discipline that evolves with advances in analytical methods and the accumulation of new data. It is established and maintained by the International Committee on Taxonomy of Viruses (ICTV), which maintains an annually updated Master Species List (MSL) of accepted taxa and names (https://ictv.global). The hierarchy for viruses includes up to 15 ranks, with realm, kingdom, phylum, class, order, family, genus, and species as the primary levels.

Previously, with VPF-Class [13], we introduced a marker-and-voting framework for taxonomic assignment and host prediction. It leveraged ≈25K pre-annotated Viral Protein Families (VPFs) from the Integrated Microbial Genomes/Virus (IMG/VR) system [11, 12] and applied consistency thresholds plus majority voting to emit lineage labels. VPF-Class remains in active use; in this work, however, we focus exclusively on taxonomic classification and do not consider host prediction further.

A single, early limitation of the first VPF-Class release was its lack of updates: it kept labels at ICTV MSL 33 and continued to use a ≈25K VPF catalogue despite the availability of a ≈227K set [2], which would enable broader marker coverage and finer taxonomic resolution. A second limitation of the first release was methodological: its architecture could not exploit the full structure of the high-dimensional marker patterns.

One possible methodological separation for viral taxonomic assignment tools distinguishes marker-driven approaches, which detect conserved protein domains and aggregate per-marker evidence into lineage calls (including, among others, VPF-Class [13], geNomad [2], and VIRGO [14]), from learning-based approaches (including, among others, ViTax [4] and PhaGCN [6]), which infer taxonomy from sequence, or network derived representations via hierarchical decision rules. We use this distinction to guide the reader and to frame how our method integrates interpretable marker signals with learned decision mechanisms.

Viral genome classification is challenging because evolutionary signal is sparse and often confined to distantly related proteins and fragmented gene repertoires. Models based on *Hidden Markov Models* (HMMs) capture conserved domains beyond pairwise alignment, while neural networks can integrate high-dimensional signals into compact representations for classification. Here we present VPF-Class 2.0, a taxonomy-centred update that combines HMM-derived marker evidence with a lightweight supervised model to produce scalable and interpretable taxonomic assignments.

## Material and Methods

This section summarizes the workflow used to convert viral genome sequences into genus/family assignments. Predicted proteins are matched to profile HMM markers, aggregated into normalized per-genome feature vectors, and classified with a lightweight supervised model consistent with the selected ICTV release. The following subsections describe the workflow overview, the neural network classifier, the marker profiles, and the datasets.

### Workflow overview

The *VPF-Class 2* pipeline converts viral genomes into family- and genus-level predictions (Fig. 1). Input sequences are processed with *prodigal-gv* [2], an adapted version of *prodigal* [5], to predict open reading frames (*ORFs*). Predicted proteins are then scanned against a collection of 227,897 HMM profiles [2]. Hits with *e-value >* 10^−3^ are discarded and the remaining matches are aggregated per genome into a marker *hit* vector. Let *v*_*i*_ denote a viral genome and *p*_*i*1_, …, *p*_*im*_ its predicted proteins. The hit vector is

**Figure 1.**
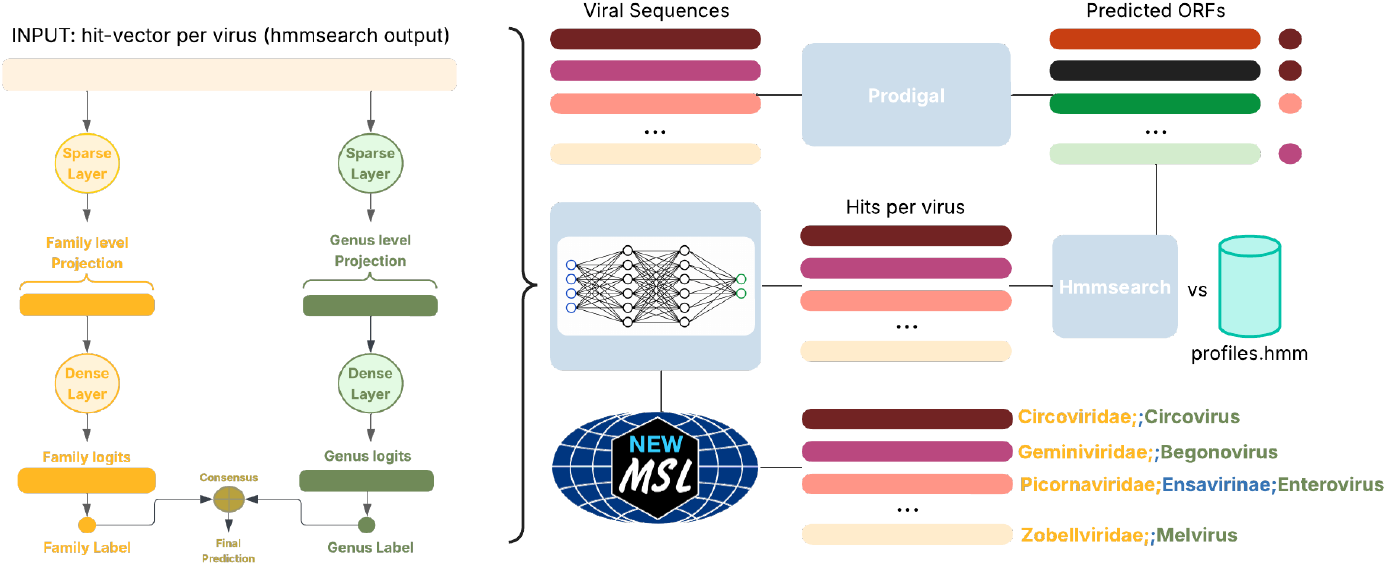
Overview of the *VPF-Class 2* workflow. Viral genomes are gene-called with prodigal, proteins are scanned with *hmmsearch* against HMM profiles, and hits are aggregated into sparse feature vectors. A sparse neural network with parallel genus- and family-level outputs produces logits that determine the final taxonomic assignment. Models are trained on ICTV MSL reference data to ensure release-consistent predictions.

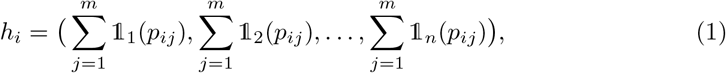

where *n* is the number of marker profiles and 𝟙_*k*_(*p*_*ij*_) = 1 if *p*_*ij*_ hits marker *k* and 0 otherwise. Because hit counts depend on genome length (Fig. 2b), we apply *ℓ*_2_ normalization,

**Figure 2.**
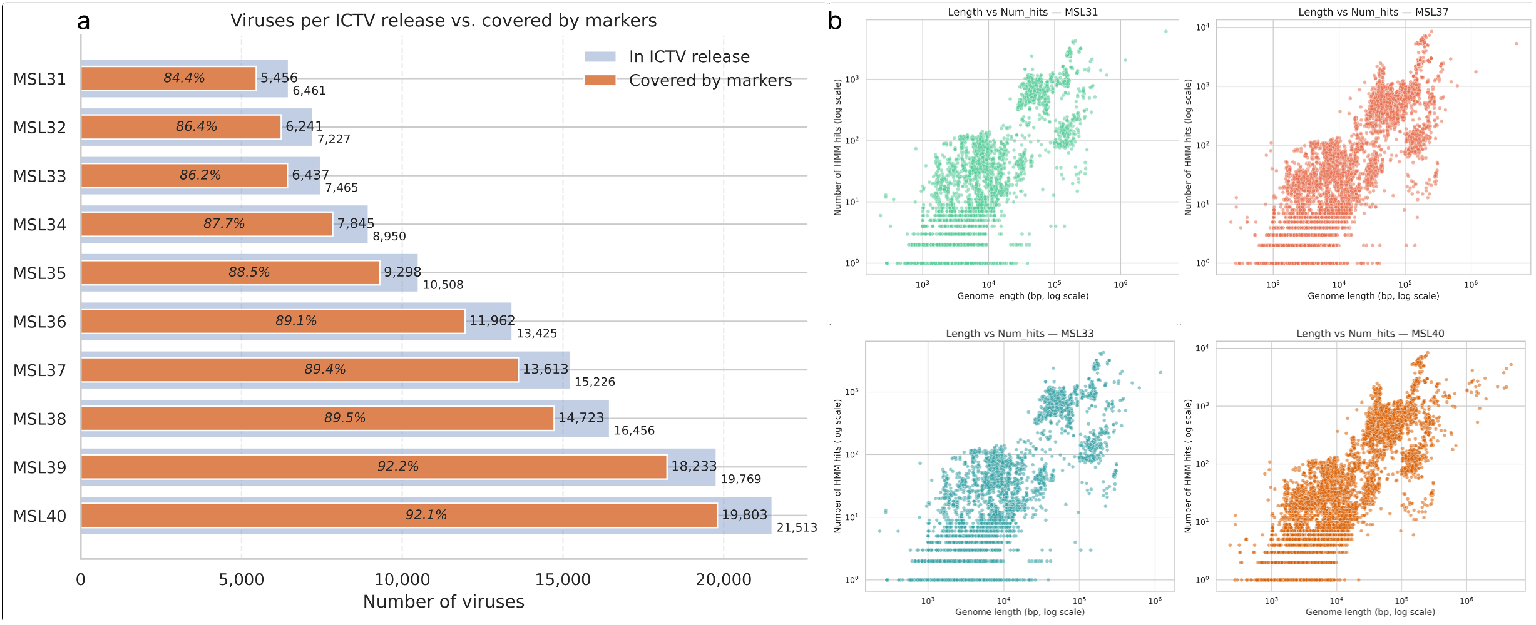
Coverage of ICTV genomes by marker profiles across releases and dependence on genome length. **(a)** Total genomes per ICTV MSL release (MSL 31–MSL 40) and the fraction with at least one marker hit (percentages indicate coverage). **(b)** Genome length versus number of HMM hits for selected releases (MSL 31, MSL 33, MSL 37, MSL 40; log scales). Each point represents one genome.

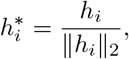

so that features reflect relative marker composition rather than absolute hit counts. The normalized vectors are then provided to the neural classifier described in the following sections.

#### Neural Network Classifier

*VPF-Class 2* uses a sparse feed-forward classifier operating on normalized marker vectors 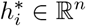, where *n* is the number of marker profiles. The network has one hidden layer of size *d* = 2024 with ReLU and dropout (*p* = 0.3), followed by a linear output layer producing logits over *C*_*g*_ genus labels:

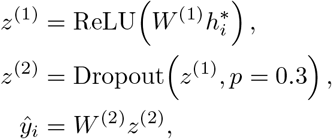

with *W* ^(1)^ ∈ ℝ^*d*×*n*^ and 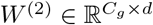. The predicted genus is arg max(*ŷ*_*i*_) and max(*ŷ*_*i*_) is used as a *confidence score*.

An analogous model is trained in parallel at the family rank, replacing *C*_*g*_ by *C*_*f*_ family labels. Genus- and family-level outputs are then combined to produce a rank-consistent lineage: genus predictions are used when sufficiently confident, otherwise the family prediction is reported (see Supplementary Section S1).

#### Marker profiles

We used the profile HMM marker collection introduced by Camargo et al. [2], which constitutes the marker set used by *geNomad* for taxonomic classification. The full catalogue contains 227,897 profiles representing protein families associated with chromosomes, plasmids, and viruses; a subset of 161,862 profiles is virus-specific.

To quantify marker coverage on ICTV references, we ran *hmmsearch* on all proteins predicted from ICTV genomes (MSL #40, release v1) using an *e*-value threshold of 10^−3^. Under this setting, 62.02 % of the virus-specific profiles were matched by at least one ICTV protein. Repeating the same analysis across earlier ICTV releases showed lower coverage, with coverage increasing with release size (Fig. 2).

By default, *VPF-Class 2* uses the 161,862 virus-specific profiles to maximize specificity, but it can optionally use the full 227,897 profile set. This option accounts for cases where viral genomes primarily match profiles outside the virus-specific subset; in MSL #40 we identified 33 such genomes, which 32 were still correctly assigned by *VPF-Class 2* (see Supplementary Table 4). Retaining both modes facilitates future retraining as ICTV releases expand and diversify.

### Datasets

We trained *VPF-Class 2* on curated ICTV reference genomes and evaluated it on large metagenomic collections without genus/family ground truth. Training used the most recent ICTV release available at the time of writing (*MSL #40, release v1* ), with genus and family labels taken from the official taxonomy. The label distribution is highly imbalanced: 52.20 % of genera (1,967/3,768) and 20.65 % of families (76/368) are represented by a single genome, which limits standard cross-validation; the evaluation strategy is described in the Results section. The same preprocessing and training protocol was applied across historical MSL releases, and final models were trained on all sequences available for each release.

To assess behaviour on real-world viromes, we analysed the Global Ocean Virome 2.0 (GOV 2.0) [3] and the Gut Phage Database (GPD) [15]. These datasets comprise hundreds of thousands of metagenome-derived viral sequences and provide a realistic setting to evaluate annotation coverage, confidence scores, and taxonomic consistency under open and highly diverse conditions.

## Results

We evaluate *VPF-Class 2* on curated ICTV benchmarks and large-scale metagenomic datasets. We first quantify accuracy and robustness across taxonomic ranks on ICTV reference data, then compare against state-of-the-art classifiers under release-matched conditions. Finally, we analyse behaviour on unlabelled viromes, focusing on annotation coverage and taxonomic consistency.

### VPF-Class 2 Performance

To obtain reproducible performance estimates under the severe class imbalance and taxonomic sparsity of the ICTV reference data, we used an adapted evaluation protocol. Standard cross-validation is not feasible because many genera are represented by a single genome and would be absent from some folds. We therefore report results under three complementary settings:

- First, genera with only one available representative were excluded from the evaluation set (1,967 out of the 3,721 annotated genera). For the remaining genus labels, we generated five independent stratified train–test splits (70–30), each using a different random seed while preserving label proportions. Across these splits, *VPF-Class 2* achieved a stable mean accuracy of 0.826 at genus level and 0.982 at family level, with limited variability across partitions (Table 1, columns *G*.*1* and *F*.*1* ).
- Secondly, genera represented by a single genome were assigned exclusively to the test partitions and excluded from training. Under this setting, the training set contains 12,485 sequences and the test set 7,318, including 1,967 sequences from genera entirely unseen during training. Consequently, 26.88 % of the test sequences correspond to unknown genus labels, reducing genus-level accuracy to a mean of 0.604 across splits (Table 1, column *G*.*2* ). These 1,967 unseen genera map to only 103 previously unseen families and represent 2.04 % of the test set; accordingly, family-level performance remains stable, with a mean accuracy of 0.948 (Table 1, column *F*.*2* ).
- Finally, we evaluated model stability when single-representative genera were incorporated into the training set. Here, the 1,967 single-genome genera were added to training, while the test partitions were kept identical to those of the first experiment. Under this setting, *VPF-Class 2* achieved a mean genus-level accuracy of 0.824 across splits, virtually identical to the baseline value of 0.826 (Table 1, column *G*.*3* ).

**Table 1.**
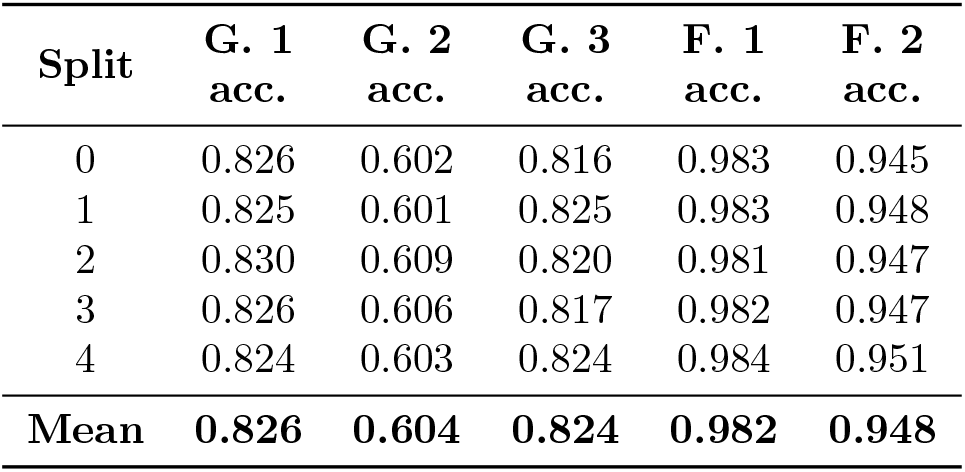
Mean classification accuracy across five stratified train–test splits under three evaluation settings. Columns *G*.*1* /*F*.*1* : baseline (single-representative genera excluded); *G*.*2* /*F*.*2* : single-representative genera included only in test; *G*.*3* : single-representative genera included in training (test identical to baseline).

| Split | G. 1<br>acc. | G. 2<br>acc. | G. 3<br>acc. | F. 1<br>acc. | F. 2<br>acc. |
| --- | --- | --- | --- | --- | --- |
| 0 | 0.826 | 0.602 | 0.816 | 0.983 | 0.945 |
| 1 | 0.825 | 0.601 | 0.825 | 0.983 | 0.948 |
| 2 | 0.830 | 0.609 | 0.820 | 0.981 | 0.947 |
| 3 | 0.826 | 0.606 | 0.817 | 0.982 | 0.947 |
| 4 | 0.824 | 0.603 | 0.824 | 0.984 | 0.951 |
| Mean | **0.826** | **0.604** | **0.824** | **0.982** | **0.948** |

Balanced accuracy and macro-F1 scores corresponding to all three experimental settings are reported in Supplementary Material S3.

#### Performance by genome properties

Genus-level performance was stratified by Baltimore genome type and by genome length. Performance is highest for dsDNA viruses, where *VPF-Class 2* reaches 0.941 accuracy (0.893 balanced accuracy; 0.879 macro-F1) over 7,155 genomes. Other genome types show more variable behaviour and smaller sample sizes (Supplementary Table 7).

A similar trend is observed with genome length, with performance improving as sequences get longer. Averaged over genomes longer than 5 kb (weighted by the number of sequences per bin; *N* = 11,495), *VPF-Class 2* reaches 0.931 accuracy, 0.849 balanced accuracy, and 0.827 macro-F1 (Supplementary Table 8).

#### Confidence threshold and annotation trade-off

In addition to the train–test evaluations, we analysed the relationship between the confidence score of *VPF-Class 2*, annotation coverage, and genus-level accuracy on the complete *MSL 40* dataset using the fully trained model. As shown in Fig. 3, increasing the confidence threshold monotonically reduces the fraction of annotated sequences while improving accuracy among retained predictions.

**Figure 3.**
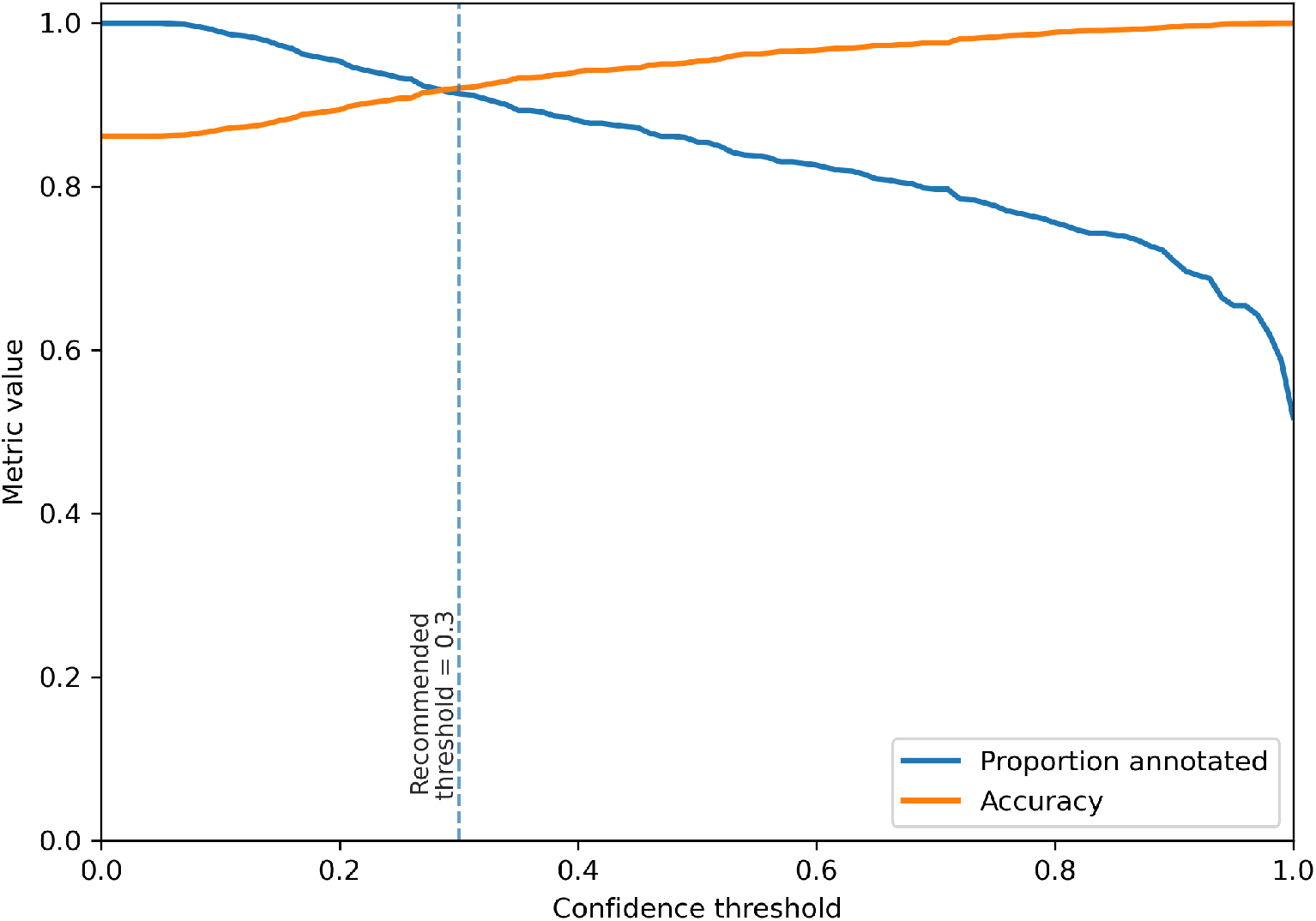
Annotation coverage (blue) and genus-level accuracy (orange) as functions of the confidence threshold. The dashed line indicates the recommended operating point at 0.3.

A threshold of 0.3 emerges as a practical operating point, retaining a large proportion of genomes while already achieving high accuracy, with only marginal improvements at more restrictive values. Accordingly, 0.3 represents a balanced compromise between annotation coverage and predictive reliability. The model produces predictions for all sequences with detectable marker hits, and the confidence score should be interpreted as a post hoc reliability indicator rather than a strict decision rule.

#### Interpretability of Taxonomic Assignments

Genus-level performance varies substantially across families, suggesting that some errors are driven by intrinsic limitations of the HMM feature space rather than by model instability. We therefore performed a post-hoc analysis to quantify how much genus-specific signal is present in the activated marker profiles within each family.

Let *F* denote a viral family and *G*_*F*_ its set of genera. For a virus *v* from genus *g* ∈ *G*_*F*_, let *H*(*v*) be the set of HMM profiles for which *v* has at least one hit (presence/absence). For each HMM profile *j* observed in family *F*, we estimate the family-conditioned probabilities

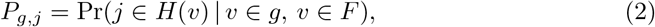

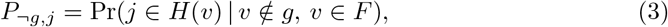

computed empirically from viruses in *F* . Families with a single genus are excluded because *P*_¬*g,j*_ is undefined.

We define a genus-specific weight for marker *j* as a smoothed log-ratio,

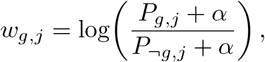

with *α >* 0 to avoid undefined values. The marker-support score of a virus *v* from genus *g* is then

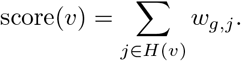

When viruses are stratified by this score, genus-level accuracy increases markedly with score (Table 2), consistent with the interpretation that low-score cases lack genus-specific marker support.^1^

**Table 2.**
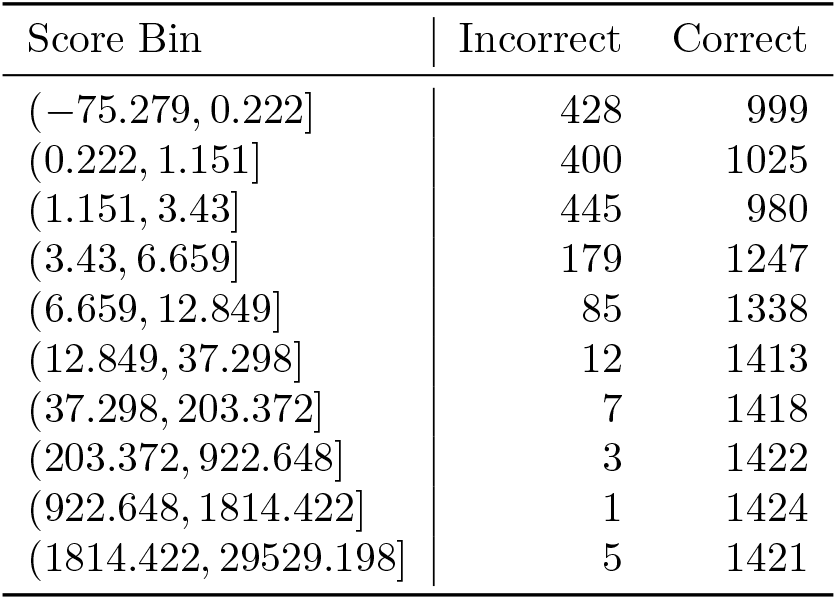
Genus-level outcomes on *MSL 38* stratified by the HMM-based score. For each score bin, we report the number of correctly and incorrectly classified viruses.

| Score Bin | Incorrect | Correct |
| --- | --- | --- |
| (−75.279, 0.222] | 428 | 999 |
| (0.222, 1.151] | 400 | 1025 |
| (1.151, 3.43] | 445 | 980 |
| (3.43, 6.659] | 179 | 1247 |
| (6.659, 12.849] | 85 | 1338 |
| (12.849, 37.298] | 12 | 1413 |
| (37.298, 203.372] | 7 | 1418 |
| (203.372, 922.648] | 3 | 1422 |
| (922.648, 1814.422] | 1 | 1424 |
| (1814.422, 29529.198] | 5 | 1421 |

To summarise ambiguity at the family level, let *S*_*F*_ be the set of HMM profiles observed in *F* and represent each genus *g* ∈ *G*_*F*_ by 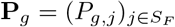. We define the intrinsic genus-level difficulty of family *F* as the mean pairwise cosine similarity between these genus profiles,

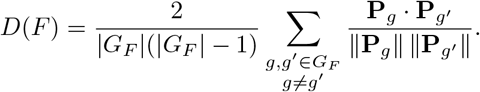

Higher *D*(*F* ) indicates stronger overlap in marker usage across genera and is associated with lower genus-level accuracy (Fig. 4).

**Figure 4.**
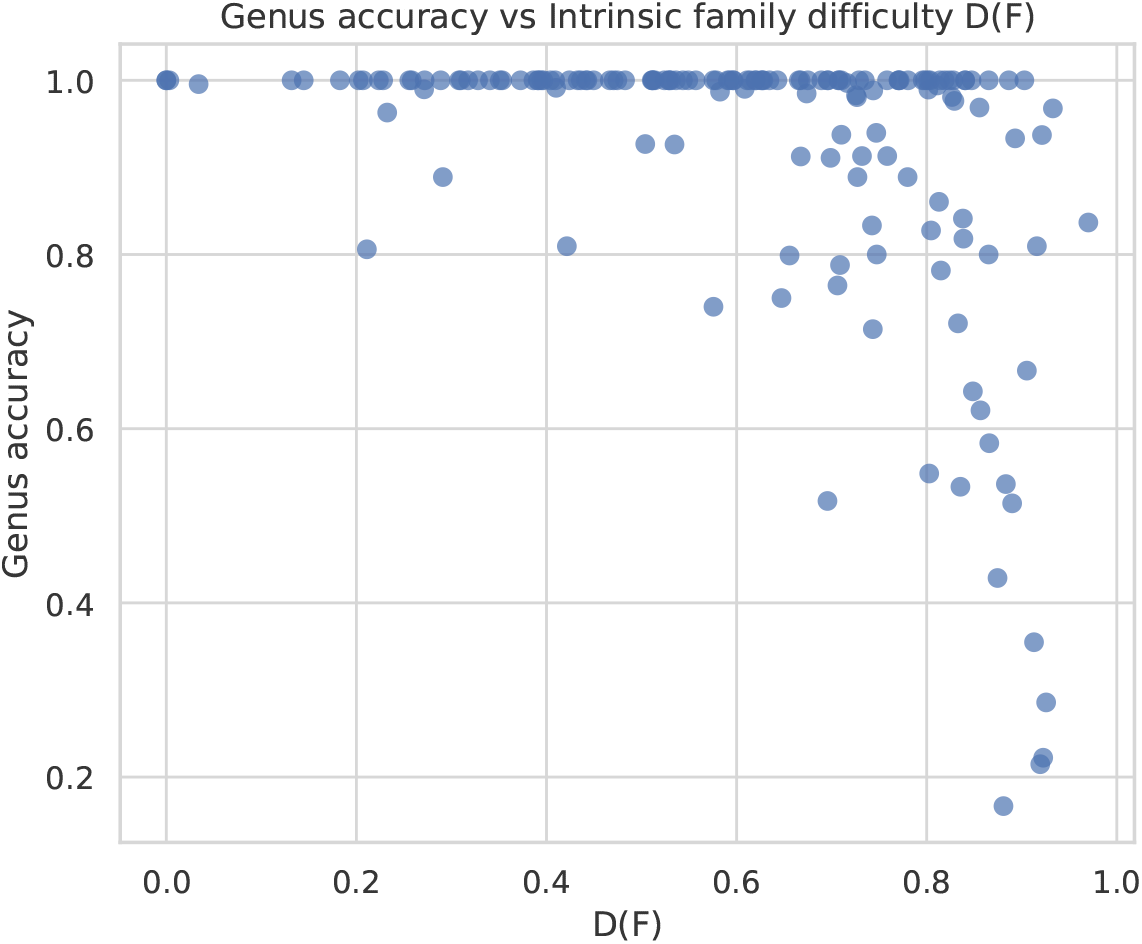
Genus-level classification accuracy as a function of intrinsic family difficulty *D*(*F* ). Higher values of *D*(*F* ), reflecting increased similarity between genus-level HMM profiles within a family, are associated with lower classification accuracy.

### Comparative performance VPF-Class 2

Comparing viral taxonomic classification tools is challenging, as each method is typically trained and evaluated on reference datasets corresponding to specific ICTV releases. Differences in taxonomic coverage and reference completeness can substantially affect reported performance, complicating direct comparisons. To ensure fairness, *VPF-Class 2* was retrained on datasets aligned with the ICTV releases and taxonomic scopes used by each competing method whenever this information was available.

Unless otherwise specified, comparisons were performed using the version of *VPF-Class 2* trained on all ICTV sequences available for the corresponding release.

When other tools were trained on restricted taxonomic subsets, equivalent subsets were used to retrain or evaluate *VPF-Class 2*, ensuring matched evaluation conditions.

#### VPF-Class, first version

The first version of the *VPF-Class* tool [13], relied on a rule-based classification approach built upon the same general pipeline: protein prediction from viral genomes followed by profile-based domain detection using *hmmsearch*. Taxonomic assignment was then performed through a voting scheme that combined evidence from different types of detected markers.

To assess the improvements introduced in *VFP-Class 2*, we performed a series of comparative experiments against the original *VPF-Class* pipeline. These analyses were designed to isolate the contribution of the methodological enhancements under different annotation and training conditions.

##### General Experiments Settings

Across all experiments, performance was evaluated consistently under three complementary scenarios. First, a *global* evaluation in which unannotated sequences were counted as classification errors, providing a conservative estimate that jointly reflects prediction accuracy and annotation coverage. Second, an *only annotated* evaluation, where performance was computed exclusively over the subset of sequences for which each tool produced a taxonomic assignment, thereby assessing accuracy conditional on successful annotation. Finally, an *intersection* evaluation restricted to the set of sequences annotated by both methods, enabling a direct comparison of taxonomic agreement under matched annotation conditions.

##### Experiment 1: Comparison on ICTV MSL 33 dataset

Both tools were evaluated on the ICTV MSL 33 release, the primary reference and benchmarking dataset of the original *VPF-Class*, enabling direct comparison under identical taxonomic conditions.

Under this setting, *VPF-Class 2* achieved accuracy, balanced accuracy, and macro-F1 scores of 0.79, substantially outperforming the original *VPF-Class*, which reached 0.26, 0.27, and 0.19, respectively (Fig. 5, panel a.1). In the *only annotated* scenario, accuracy increased to 0.92 for *VPF-Class 2* versus 0.47 for *VPF-Class* (Fig. 5, panel a.2), while the intersection analysis yielded 0.97 and 0.49, respectively (Fig. 5, panel a.3).

**Figure 5.**
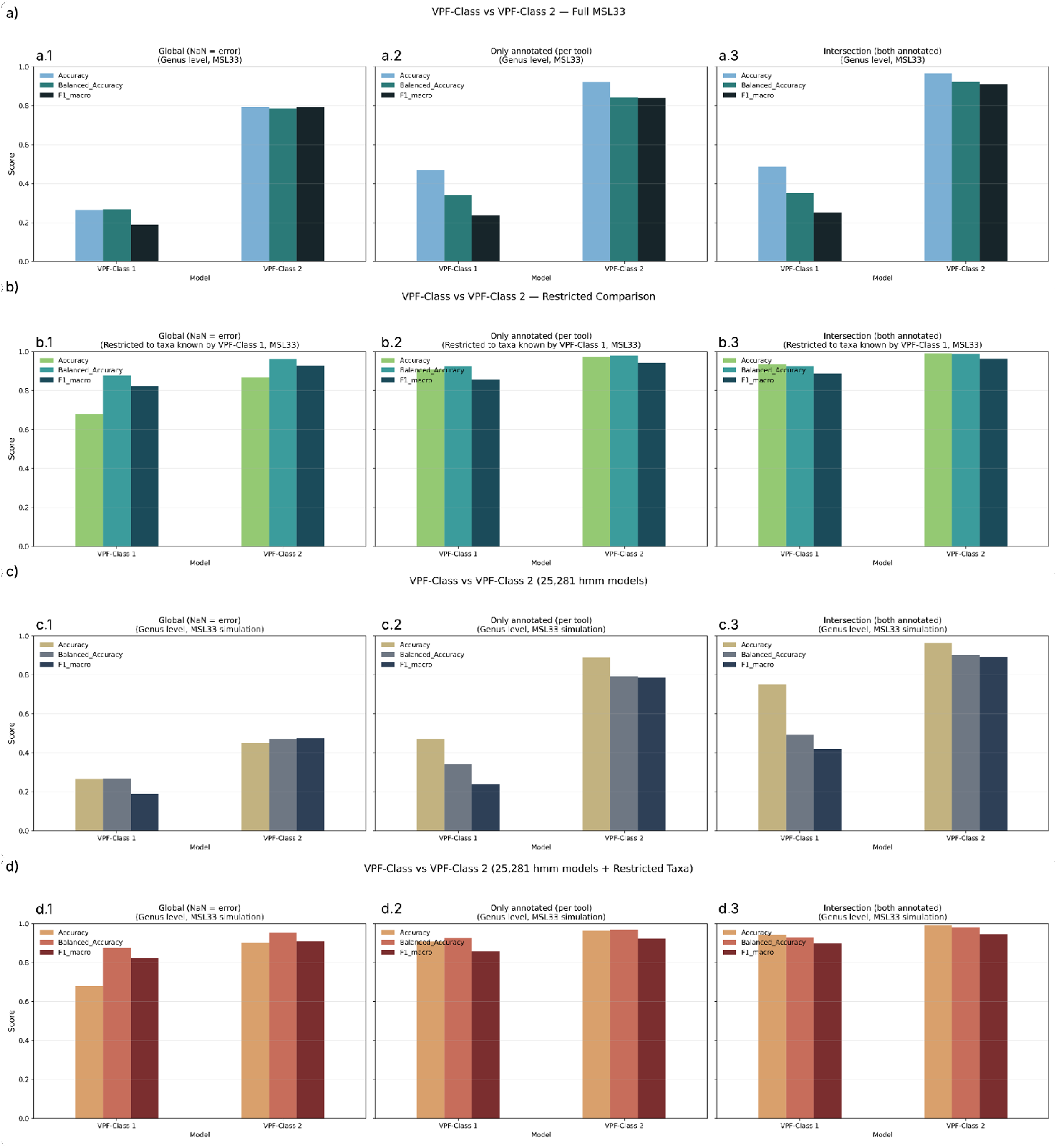
Genus-level performance comparison between *VPF-Class* (v1) and *VPF-Class 2* on the ICTV *MSL 33* dataset. Panels (a)–(d) correspond to Experiments 1–4, respectively. Columns show *global evaluation, only annotated*, and *intersection* analyses. Bars represent accuracy, balanced accuracy, and macro-F1.

##### Experiment 2: Comparison restricted to the genera predictable by *VPF-Class*

Due to its rule-based voting procedure, the original *VPF-Class* could assign only 287 of the 842 genera annotated in MSL 33. We therefore restricted the evaluation to this subset to assess performance under comparable taxonomic coverage.

In this scenario, *VPF-Class* improved to an accuracy of 0.68, balanced accuracy of 0.88, and macro-F1 of 0.82. Nevertheless, *VPF-Class 2* achieved higher scores, with 0.87, 0.96, and 0.93, respectively (Fig. 5, panel b.1). In the *only annotated* setting, accuracies were 0.97 for *VPF-Class 2* and 0.91 for *VPF-Class*, while the intersection analysis yielded 0.99 and 0.93 (Fig. 5, panels b.2, b.3).

##### Experiment 3: Retrospective comparison using the original marker set

To evaluate whether the improvements introduced by *VPF-Class 2* arise solely from the expanded marker database or from the change in classification strategy itself, a retrospective experiment was performed, where *VPF-Class 2* was retrained using the original set of ≈25,000 HMM profiles employed by *VPF-Class*.

Under these constrained conditions, the learning-based model achieved markedly higher performance, reaching an accuracy of about 0.45, a balanced accuracy of 0.47, and a macro-F1 of 0.47, whereas the original tool reached only 0.26, 0.27, and 0.19, respectively (Fig. 5, panel c.1). Across the remaining evaluation scenarios: accuracies increased to 0.89 versus 0.47 in the *only annotated* setting, and to 0.96 versus 0.75 in the intersection analysis for *VPF-Class 2* and *VPF-Class*, respectively (Fig. 5, panels c.2, c.3).

##### Experiment 4: Restricted taxonomy using the original marker set

Finally, the two previous constraints are combined: the original set of ≈25,000 marker profiles and the subset of 287 genera assignable by *VPF-Class*, defining the most restrictive evaluation scenario.

Even under these conditions, *VPF-Class 2* maintained a clear advantage, achieving accuracy, balanced accuracy, and macro-F1 scores of approximately 0.90, 0.95, and 0.91, compared to 0.68, 0.88, and 0.82 for *VPF-Class* (Fig. 5, d.1). In the *only annotated* setting, accuracies increased to 0.96 and 0.91, respectively, while the intersection analysis yielded 0.99 and 0.94 (Fig. 5, d.2, d.3).

Across all four evaluation settings, *VPF-Class 2* consistently and substantially outperformed its predecessor. Importantly, this advantage persisted even when using the original marker set and restricting the taxonomy to the genera assignable by *VPF-Class*. These results indicate that the primary source of improvement lies in the transition from a rule-based voting scheme to a supervised learning framework, rather than in expanded marker coverage alone.

#### geNomad

*geNomad* [2] is one of the most widely used and up-to-date frameworks for viral discovery and classification. It relies on the same set of ≈227K marker profiles that *VPF-Class 2* builds upon, but its classification procedure is based on a homology-driven, rule-based hierarchy that assigns taxonomy by aggregating marker evidence according to predefined thresholds.

##### Comparison with geNomad

*geNomad* currently provides viral annotations consistent with the ICTV *MSL 39* release. Accordingly, the evaluation was conducted on sequences from this release to ensure comparability between both tools. As *geNomad* produces taxonomic assignments only up to the family level, the comparison was limited to this rank and followed the same three evaluation schemes used in previous experiments (Fig. 6, top panels). When considering all sequences, *VPF-Class 2* achieved notably higher performance, with an accuracy of approximately 0.91 compared to 0.59 for *geNomad*. The same trend was observed for balanced accuracy (0.85 vs. 0.41) and macro-F1 (0.87 vs. 0.43), reflecting a substantial improvement in overall predictive reliability. In the *only annotated* and *intersection* evaluations, both tools exhibited high-performance, achieving accuracy scores of around 0.99 for *VPF-Class 2* and 0.91 for *geNomad*. However, the higher global performance of *VPF-Class 2* indicates increased sensitivity and broader annotation coverage.

**Figure 6.**
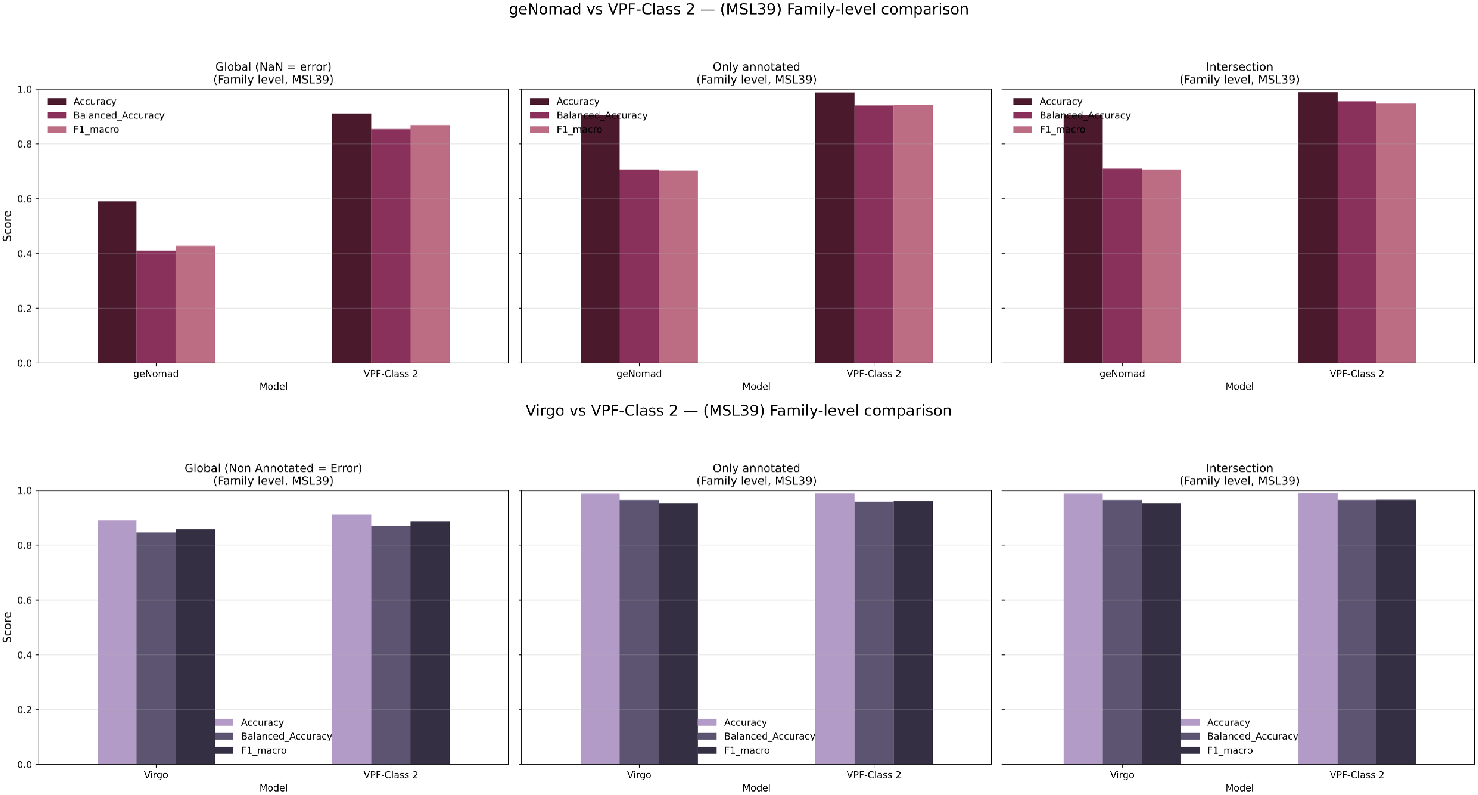
Family-level performance comparison of *VPF-Class 2* with *geNomad* (top) and *VIRGO* (bottom) on the ICTV *MSL 39* dataset. Columns show global, *only annotated*, and intersection evaluations. Bars represent accuracy, balanced accuracy, and macro-F1.

#### VIRGO

*VIRGO* [14] is a recent viral taxonomy framework that, like *VPF-Class 2*, relies on the set of *geNomad* viral marker profiles. In contrast to a learning-based approach, *VIRGO* performs classification through a similarity-driven procedure that quantifies the overlap between marker subsets using a bidirectional subsethood metric.

##### Comparison with VIRGO

The evaluation was conducted using sequences from the ICTV *MSL 39* release, restricting predictions to the family level to ensure methodological consistency between tools (Fig. 6, bottom panels). When all sequences were considered, including those left unannotated by either method, *VPF-Class 2* achieved slightly higher performance than *VIRGO*, with accuracy values around 0.91 versus 0.89, alongside marginal improvements in balanced accuracy and macro-F1. In this global setting, *VPF-Class 2* left 1,536 sequences without family-level assignment, whereas *VIRGO* did not annotate 1,938 sequences.

In the *only annotated* and *intersection* scenarios, both tools reached very high accuracy values, close to 0.99, reflecting reliable family-level assignments whenever sufficient marker evidence is available. In these restricted evaluations, performance differences between the two methods were minor, with *VPF-Class 2* matching or slightly exceeding *VIRGO* across most metrics.

These results show that *VPF-Class 2* achieves family-level classification performance comparable to, and in some cases marginally better than, *VIRGO*, while additionally providing genus-level predictions and annotating a larger fraction of sequences.

#### ViTax

*ViTaX* adopts a fundamentally different paradigm for viral taxonomic classification by framing genomic sequences as a language modelling problem [4]. Built upon *HyenaDNA*, a foundation model for long-range genomic sequences at single-nucleotide resolution, it learns latent representations directly from raw nucleotide sequences without relying on curated protein-domain markers or homology-based features. To mitigate taxonomic imbalance, the method incorporates supervised prototypical contrastive learning, while a belief-mapping tree based on the Lowest Common Ancestor algorithm enables assignment to the lowest confident taxonomic rank in open-set scenarios.

##### Comparison with ViTaX

For each input sequence, *ViTaX* may produce a genus- or family-level assignment, return an *unclassified* label, or assign it to a higher taxonomic rank when the predicted genus is considered unknown. Accordingly, the definition of classification error depends on how LCA and *unclassified* outputs are treated. All comparisons were performed using the ICTV *MSL 38* taxonomy, consistent with the reference framework adopted by *ViTaX*.

As *ViTaX* was trained to recognize a fixed set of 631 genera, the evaluation strategy differs from previous comparisons. Results are therefore reported for the sequences annotated by each tool individually and for their intersection, since a global evaluation would disproportionately penalize *ViTaX* outside its intended taxonomic scope.

Within this framework, three complementary experiments were conducted. First, both tools were evaluated on the full MSL 38 dataset, considering only explicit genus- or family-level assignments as valid annotations and excluding LCA and *unclassified* outputs. Second, the analysis was restricted to sequences belonging to the 631 genera covered by *ViTaX*, excluding LCA predictions from the comparison. Finally, a more stringent evaluation was performed on the same restricted subset, in which both LCA and *unclassified* outputs were compared against the ground truth and counted as errors. Together, these experiments capture different operational interpretations of open-set predictions and allow assessment of the relative behaviour of both methods.

- On the complete ICTV *MSL 38* dataset (Fig. 7, top panels), we evaluated only explicit genus/family assignments, excluding LCA and *unclassified* outputs. At family level, *VPF-Class 2* achieved near-perfect performance (accuracy, balanced accuracy and macro-F1 around 0.99, 0.95 and 0.95, respectively), whereas *ViTaX* reached values around 0.63, 0.38 and 0.33, respectively. At genus level, *VPF-Class 2* obtained 0.89 accuracy and 0.77 macro-F1, compared to 0.64 and 0.46 for *ViTaX*. The same gap persisted on the intersection of annotated sequences (family: 0.99 vs 0.65 accuracy; genus: 0.93 vs 0.67 accuracy; macro-F1 0.85 vs 0.48). Family- and genus-level metrics are computed on different subsets because excluding LCA removes more sequences at genus rank.
- To reflect *ViTaX*’s intended operating regime, we restricted the evaluation to sequences whose true labels belong to the 631 genera used by *ViTaX* (Fig. 7, middle panels), again excluding LCA and *unclassified*. Under this closed-set setting, both tools achieved near-perfect family-level performance (both close to 1.00). At genus level, *ViTaX* was slightly better (accuracy around 0.99) than *VPF-Class 2* (accuracy around 0.97), with consistent trends for balanced accuracy and macro-F1. Similar patterns were observed on the intersection subset.
- Finally, under the same restriction but counting LCA outputs as errors (Fig. 7, bottom panels), *VPF-Class 2* retained perfect family-level performance (= 1.00), while *ViTaX* decreased slightly (family accuracy of 0.98 on annotated predictions; 0.99 on the intersection; macro-F1 0.97). At genus level, both methods remained high and close: *VPF-Class 2* reached 0.96 accuracy (0.97 balanced accuracy; 0.97 macro-F1) versus 0.94 (0.94; 0.96) for *ViTaX*, with similar results on the intersection.

**Figure 7.**
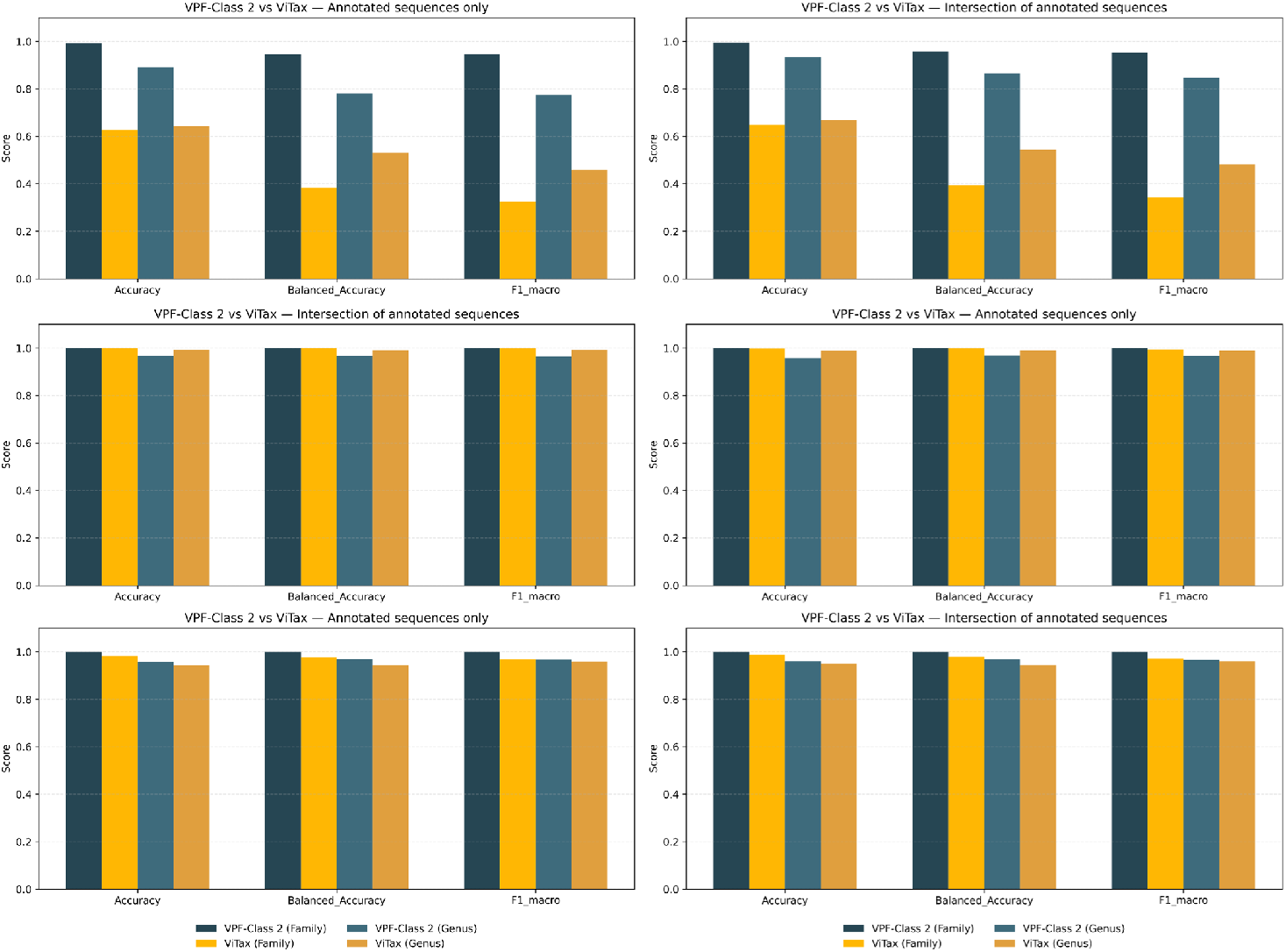
Comparison between *VPF-Class 2* and *ViTaX* on the ICTV *MSL 38* dataset. Rows (top to bottom) correspond to the three evaluation scenarios described in the main text. Columns show annotated-only and intersection analyses for both family- and genus-level performance.

Overall, these results highlight the distinct design philosophies of the two approaches. *ViTaX* is optimized for open-set classification, favouring conservative LCA assignments when confidence is low, whereas *VPF-Class 2* aims to provide taxonomic predictions without an explicit open-set mechanism. When evaluated within a shared closed-set taxonomy and penalizing conservative outputs, *VPF-Class 2* matches or slightly exceeds the performance of *ViTaX* at both family and genus levels.

#### PhaGCN3

*PhaGCN* is one of the most advanced graph-based frameworks for viral taxonomic classification. It represents viral genomes as nodes in a gene-sharing network, with edges encoding shared protein clusters between sequences, and applies graph convolutional networks to propagate taxonomic information across the resulting topology.

##### Comparison with PhaGCN

To compare *VPF-Class 2* with *PhaGCN* on the ICTV *MSL 38* (v3), we evaluated both methods under three scenarios designed to account for their fundamentally different annotation strategies.

First, we considered the complete dataset, where all sequences were included regardless of marker coverage. In this setting, *PhaGCN* achieved higher overall performance, reflecting its ability to classify a larger fraction of sequences. While *VPF-Class 2* reached an accuracy close to 0.80, with balanced accuracy and macro-F1 around 0.76, *PhaGCN* attained values above 0.90 across all metrics (Fig 8).

**Figure 8.**
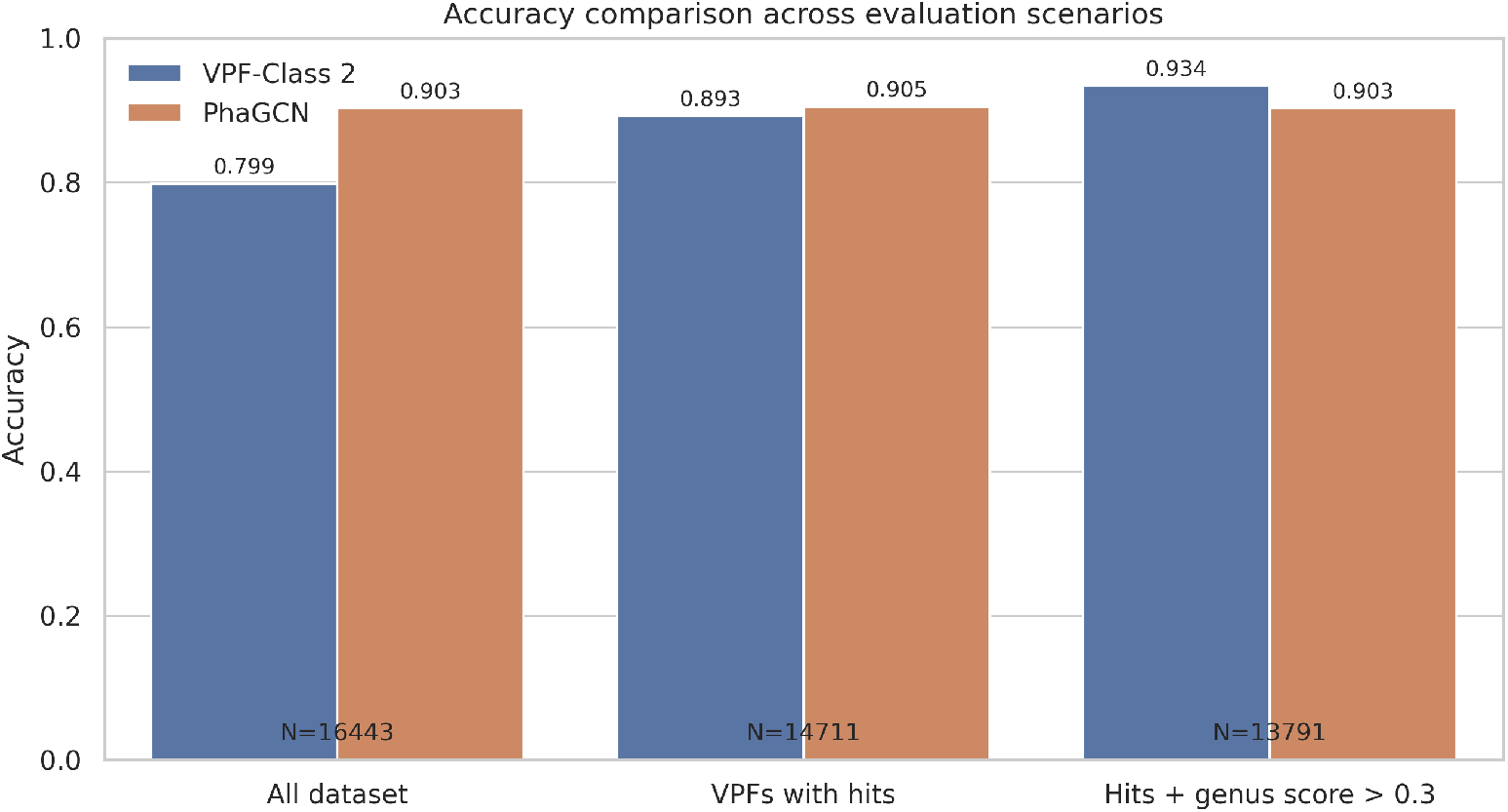
Genus-level performance of *VPF-Class 2* and *PhaGCN* on the ICTV *MSL 38* release (v3).

In a second evaluation, the comparison was restricted to sequences for which *VPF-Class 2* produced at least one marker hit. Under this condition, the performance gap between both methods narrowed substantially. In terms of accuracy, both approaches converged to comparable values close to 0.9, indicating that, when marker evidence is available, *VPF-Class 2* can achieve a level of predictive correctness similar to that of *PhaGCN*. However, *PhaGCN* retained an advantage in balanced accuracy and macro-F1, with both metrics remaining close to 0.9, whereas *VPF-Class 2* reached values around 0.78 and 0.77, respectively.

Finally, we considered a more restrictive evaluation focused on high-confidence predictions produced by *VPF-Class 2*. In this setting, only sequences with a genus-level confidence score greater than 0.3 were retained, which still accounted for more than 83 % of the viral sequences in the ICTV *MSL 38* release. Within this subset, *VPF-Class 2* achieved an accuracy of 0.93, together with balanced accuracy and macro-F1 values around to 0.90. Under the same conditions, *PhaGCN* reached a slightly lower accuracy of 0.90, while exhibiting nearly identical balanced accuracy and macro-F1 scores than *VPF-Class 2*.

These results highlight complementary strengths between the two approaches. *PhaGCN* benefits from broader annotation coverage and more uniform performance across taxa, whereas *VPF-Class 2* achieves comparable performance when marker evidence is available and matches or slightly exceeds *PhaGCN* in high-confidence regimes. This indicates that the marker-based learning framework can deliver highly reliable taxonomic assignments when sufficient signal is present, while remaining more sensitive to class imbalance in low-evidence scenarios.

Notably, length-dependent differences were also observed when comparing both methods. Using a 7 kb threshold to partition the dataset (Table 3) into two approximately equal-sized groups, clear length-dependent differences emerged. For genomes shorter than 7 kb, *PhaGCN* substantially outperformed *VPF-Class 2*, achieving an accuracy close to 0.95 compared to approximately 0.81 for *VPF-Class 2*. In contrast, for genomes longer than 7 kb, *VPF-Class 2* surpassed *PhaGCN*, reaching an accuracy of about 0.96 versus 0.89, respectively.

**Table 3.** Accuracy of *VPF-Class 2* and *PhaGCN* stratified by genome length (≤7 kb vs *>*7 kb) on sequences predicted by both tools.

| Length Bin | N | VPF-Class 2 | PhaGCN |
| --- | --- | --- | --- |
| 0–7000 | 6762 | 0.814 | 0.948 |
| $> 7000$ | 7735 | 0.959 | 0.892 |

This behaviour is consistent with the genome-type and length stratifications described above. In the ICTV reference dataset, longer genomes are predominantly associated with double-stranded DNA viruses, which also represent the group where *VPF-Class 2* shows the most stable and highest performance (Supplementary Fig. 11). Conversely, shorter genomes are more frequent among single-stranded RNA and small DNA viruses, where *PhaGCN* classification remains more robust.

### Real datasets

We evaluated *VPF-Class 2* on GOV 2.0 and GPD datasets without curated genus/family labels, focusing on coverage, rank consistency, and overlap with established tools.

#### Global Ocean Virome 2.0

We analysed the GOV 2.0 dataset filtered to contigs ≥5 kb or circular, comprising 488,131 sequences (195,728 *>*10 kb). *VPF-Class 2* detected at least one HMM hit for most sequences; 3,620 contigs lacked marker hits (515 among those *>*10 kb). Applying a genus-level confidence threshold of 0.3 yielded confident annotations for ∼11% of all sequences and ∼15% of those *>*10 kb.

Among confident predictions, assignments were dominated by *Caudoviricetes* (92.65%). Smaller fractions of predictions were assigned to *Megaviricetes* (1.31%), *Arfiviricetes* (1.18%), and *Virophaviricetes* (0.52%). At the family level, 40.65% of predictions mapped to genera without an ICTV-recognized family; among classified families, *Pachyviridae* (14.55%), *Pervagoviridae* (7.99%), and *Salasmaviridae* (4.10%) were most frequent. Genus-level predictions were enriched in *Callevirus* (14.06%), *Inhavirus* (12.18%), *Harambevirus* (5.86%), and *Bacelvirus* (4.81%).

For comparison, we applied the same *VPF-Class 2* model trained on ICTV MSL 38 alongside *PhaGCN*. Here, *PhaGCN* produced more genus-level annotations, while *VPF-Class 2* retained a smaller high-confidence set (threshold 0.3). On the intersection, 18,236 sequences were annotated by both tools at genus level, and 10,951 (about 60%) received the same genus. The predominant family-level assignments were also consistent across tools, with *Kyanoviridae* and *Autographiviridae* among the most frequent calls.

#### Gut Phage Database

We applied *VPF-Class 2* to the Gut Phage Database (142,809 sequences). Marker coverage was high: only 281 sequences lacked HMM hits. Using a genus-level confidence threshold of 0.3, we retained 44,378 sequences (about 31%) for downstream analyses.

Confident predictions were dominated by *Caudoviricetes* (nearly 97%). At the family level, 52.2% of predictions mapped to genera without an ICTV-recognized family; among classified families, *Suoliviridae, Guelinviridae, Ludisviridae*, and *Intestiviridae* were most frequent.

For comparison with *PhaGCN*, we used the same *VPF-Class 2* model trained on ICTV MSL 38. After applying the same confidence threshold, *VPF-Class 2* produced 42,018 genus-level annotations, whereas *PhaGCN* annotated 21,515 sequences; 13,318 sequences were annotated by both tools (Fig. 9). Within this intersection, 11,860 sequences (89.05%) received the same genus. Among disagreements, 89.6% still agreed at the family level, and only about 1% of the intersection corresponded to different family assignments.

**Figure 9.**
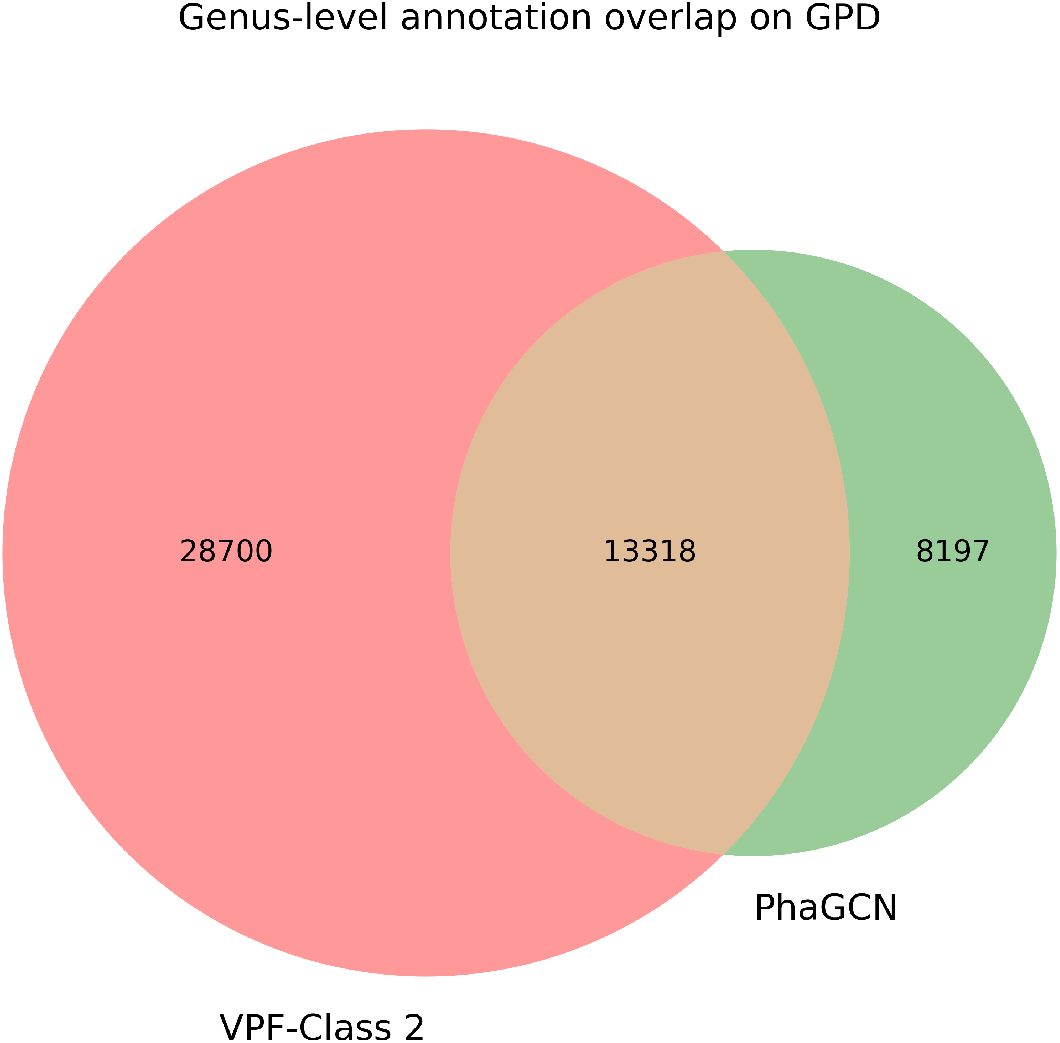
Genus-level annotation overlap between *VPF-Class 2* and *PhaGCN* on the Gut Phage Database (GPD). *VPF-Class 2* annotations were filtered using a genus-level confidence threshold of 0.3. Numbers indicate the count of viral sequences annotated exclusively by each tool and those jointly annotated by both methods.

Together, these results indicate that, *VPF-Class 2* and *PhaGCN* converge on highly consistent taxonomic assignments for confidently annotated gut phage sequences, while differing primarily in the breadth of their genus-level coverage.

## Conclusions

In this work, we presented *VPF-Class 2*.*0*, a taxonomy-centered update that markedly improves upon the original *VPF-Class* and achieves state-of-the-art performance within the marker-driven paradigm. By retaining interpretable HMM evidence while replacing rule-based voting with a lightweight supervised classifier, *VPF-Class 2*.*0* exploits high-dimensional marker dependencies that most marker-based tools do not capture. On ICTV benchmarks, it delivers near-perfect family-level performance (mean accuracy 0.982) and strong genus-level accuracy (0.826), and remains robust under extreme sparsity, with family accuracy of 0.948 even when many genera are unseen. In release-matched comparisons, it consistently outperforms *VPF-Class* v1 and surpasses representative marker-driven tools in global evaluations by combining high accuracy with broader effective annotation coverage; within shared taxonomic scope, it matches or improves upon learning- and graph-based approaches in high-confidence regimes (threshold 0.3). Finally, we introduced an interpretability layer linking genus errors to within-family marker overlap and demonstrated real-world applicability on large viromes, with high concordance against *PhaGCN* on GPD (89.05% genus agreement on the intersection) and substantial agreement on GOV 2.0 (about 60%).

## Competing interests

No competing interest is declared.

## Author contributions statement

L.V.: Methodology, software, validation, formal analysis, data curation, writing original draft, visualization. J.C.P.: Conceptualization, methodology, validation, formal analysis, writing original draft, writing—review & editing, supervision, project administration. M.L.: Supervision, resources, writing—review & editing. M.B.F: Resources, writing—review & editing. N.K.: Supervision, resources, writing—review & editing. All authors read and approved the final manuscript.

## Acknowledgments

LV, JCP and ML were supported in part from Grant PID2021-126114NB-C44 funded by MCIN/AEI/ 10.13039/501100011033. MBF and NCK were supported by the US DOE JGI (https://ror.org/04xm1d337), a DOE Office of Science User Facility, supported by the Office of Science of the US DOE operated under contract no. DE-AC02-05CH11231.”

## Supporting Information

### S1 Lineage Reconstruction and Consistency Corrections

Family- and genus-level predictions are obtained from two independently trained neural networks. Although hierarchical architectures were explored during development, they did not lead to improved predictive performance. Consequently, both taxonomic levels are inferred independently and subsequently combined during lineage reconstruction.

While this design maximizes predictive flexibility, it may lead to occasional inconsistencies between genus- and family-level predictions. In particular, the predicted genus may belong to a different family than the one independently predicted at the family level. Additionally, the family-level classifier includes a dedicated *Unknown* label, which corresponds to genomes whose annotated genus does not have an associated family in the selected ICTV release.

To provide taxonomic consistency in the reconstructed lineage, two post-processing corrections are applied (Figure 10).

**Figure 10.**
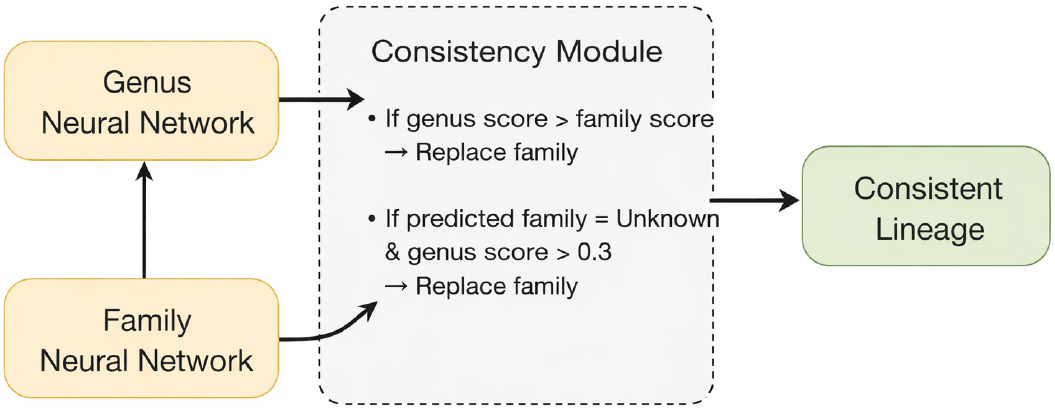
Lineage reconstruction workflow. Genus- and family-level classifiers are trained independently. A consistency module integrates both predictions by applying two post-processing rules: (i) if the genus confidence score exceeds the family confidence score, the family prediction is replaced by the family associated with the predicted genus; (ii) if the predicted family is *Unknown* and the genus confidence exceeds 0.3, the family is replaced by the genus-inferred family. The resulting lineage is thus taxonomically coherent while preserving independent model training.

#### S1.1 Score-Based Family Adjustment

In rare cases, the confidence score (maximum logit) of the genus-level prediction exceeds the confidence score of the family-level prediction for the same genome. When this occurs, the predicted family is replaced by the family induced by the predicted genus, according to the ICTV release used for training.

This correction is applied conservatively and affects only a small number of sequences. It reflects the assumption that, when the genus-level classifier expresses higher confidence than the family-level classifier, the genus prediction provides more reliable taxonomic information.

#### S1.2 Correction of Overconfident *Unknown* Family Assignments

In real dataset analyses, we observed that the family-level classifier tends to overestimate the confidence of the *Unknown* label. This behaviour can lead to inconsistencies where a genome is assigned a valid genus with a known family association, yet its family prediction is *Unknown*.

To mitigate this effect, an additional correction is applied: if the predicted family is *Unknown* and the genus-level confidence score exceeds 0.3, the family prediction is replaced by the family associated with the predicted genus.

This threshold was selected empirically to balance correction strength and robustness. Interestingly, when applying this procedure to the MSL39 dataset, no changes were observed. All genomes predicted as *Unknown* at the family level corresponded to genera that indeed lack an associated family in that ICTV release. This observation suggests that the correction mechanism primarily addresses specific dataset-dependent inconsistencies rather than introducing systematic bias.

It is important to note that this post-processing strategy does not enforce strict hierarchical consistency in all cases. In particular, when the genus-level prediction has low confidence and does not satisfy the correction criteria described above, disagreements between genus and family predictions are not overridden. In such cases, lineage reconstruction follows the family-level prediction, even if the predicted genus would correspond to a different family under the ICTV taxonomy.

This design choice reflects a conservative strategy: only sufficiently confident genus-level predictions are allowed to influence higher-level assignments. Consequently, potential inconsistencies at low genus confidence are preserved rather than forcibly reconciled, preventing the introduction of overcorrections in uncertain scenarios.

### S2 Use of non virus-specific marker profiles

By default, *VPF-Class 2* restricts feature construction to the 161,862 virus-specific HMM profiles in order to maximise taxonomic specificity and reduce noise from broadly conserved domains. However, the full collection of 227,897 profiles can optionally be used when required. In the ICTV *MSL 40* (v1) dataset, we identified 33 genomes whose detected HMM hits fell exclusively outside the virus-specific subset. As a consequence, these sequences cannot be classified by the default virus-only model, since no input features are generated. When the full marker collection is enabled, all but one of these genomes were correctly assigned at the genus level. The only misclassification corresponded to accession DQ087281.1, a *Dinovernavirus* predicted as *Cypovirus*. Notably, the predicted genus-level confidence scores for these sequences were consistently high (Table 4), indicating that non virus-specific markers can still provide strong and biologically informative signals for certain viral taxa. Retaining both operational modes therefore offers flexibility for future ICTV releases, where novel or atypical viral genomes may predominantly match profiles outside the curated virus-specific subset.

**Table 4.** Viral genus predictions and confidence scores.

| Accession | pred_genus | score_genus |
| --- | --- | --- |
| AB126195.1 | Gammainfluenzavirus | 0.917 |
| AC007209.6 | Metavirus | 0.989 |
| AF291689.1 | Cypovirus | 0.913 |
| AF389457.1 | Cypovirus | 0.913 |
| AF389467.1 | Cypovirus | 0.913 |
| AY177408.1 | Cypovirus | 0.913 |
| D83207.1 | Semotivirus | 0.900 |
| DQ077913.1 | Cypovirus | 0.913 |
| DQ087281.1 | Cypovirus | 0.913 |
| EU195327.1 | Betapartitivirus | 0.939 |
| GQ274024.1 | Nanovirus | 0.967 |
| J02207.1 | Gammaretrovirus | 0.983 |
| JN133281.1 | Nanovirus | 0.966 |
| KC588362.1 | Cypovirus | 0.913 |
| KC588370.1 | Cypovirus | 0.913 |
| KC978956.1 | Nanovirus | 0.965 |

### S3 Extended Cross-Validation Results

In the main manuscript, cross-validation experiments are summarized using overall accuracy as the primary performance metric. While accuracy provides a global estimate of predictive performance, it may be influenced by class imbalance, particularly in taxonomic classification tasks where genera and families are unevenly represented.

To provide a more comprehensive evaluation, we report here the balanced accuracy and macro-averaged F1-score for all cross-validation folds and evaluation settings. Balanced accuracy is defined as the average recall across classes and therefore accounts for class imbalance by giving equal weight to each taxon. The macro-F1 score further summarizes performance by computing the unweighted mean of per-class F1-scores, penalizing poor performance on under-represented classes.

These additional metrics allow a more nuanced interpretation of model behaviour across splits and complement the accuracy values presented in the main text.

**Table 5.** Family-level balanced accuracy and macro-F1 across cross-validation splits.

| Split | F.1 bal. acc. | F.1 macro-F1 | F.2 bal. acc. | F.2 macro-F1 |
| --- | --- | --- | --- | --- |
| 0 | 0.910 | 0.909 | 0.616 | 0.600 |
| 1 | 0.906 | 0.901 | 0.606 | 0.589 |
| 2 | 0.900 | 0.893 | 0.614 | 0.599 |
| 3 | 0.897 | 0.891 | 0.611 | 0.592 |
| 4 | 0.899 | 0.898 | 0.609 | 0.596 |
| **Mean** | **0.902** | **0.899** | **0.611** | **0.595** |

**Table 6.**
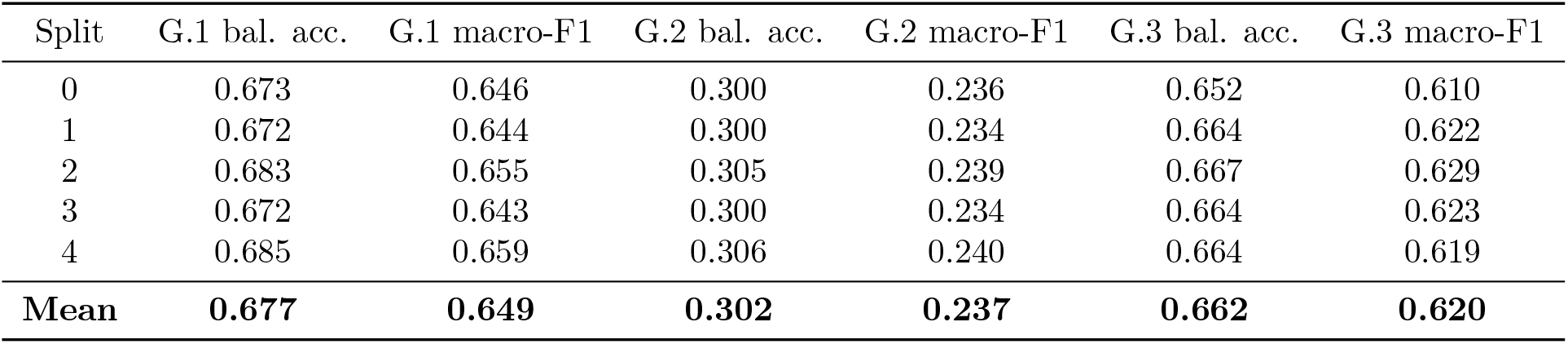
Genus-level balanced accuracy and macro-F1 across cross-validation splits under different evaluation settings.

### S4 Genome Type vs Genome Length

To characterise the structural properties of the MSL40 dataset, we examined the distribution of genome lengths across Baltimore genome types. Figure 11 shows violin plots with overlaid boxplots representing the length distributions (in base pairs, log-scale) for each genome category. The violin shapes illustrate the density of genome sizes, while the boxplots summarise the median and interquartile range, excluding extreme outliers for clarity. This analysis provides contextual information on the inherent size variability among genome types, which is relevant when interpreting downstream performance differences, as genome length directly influences the number of detectable coding regions and, consequently, the amount of functional signal available for classification.

**Figure 11.**
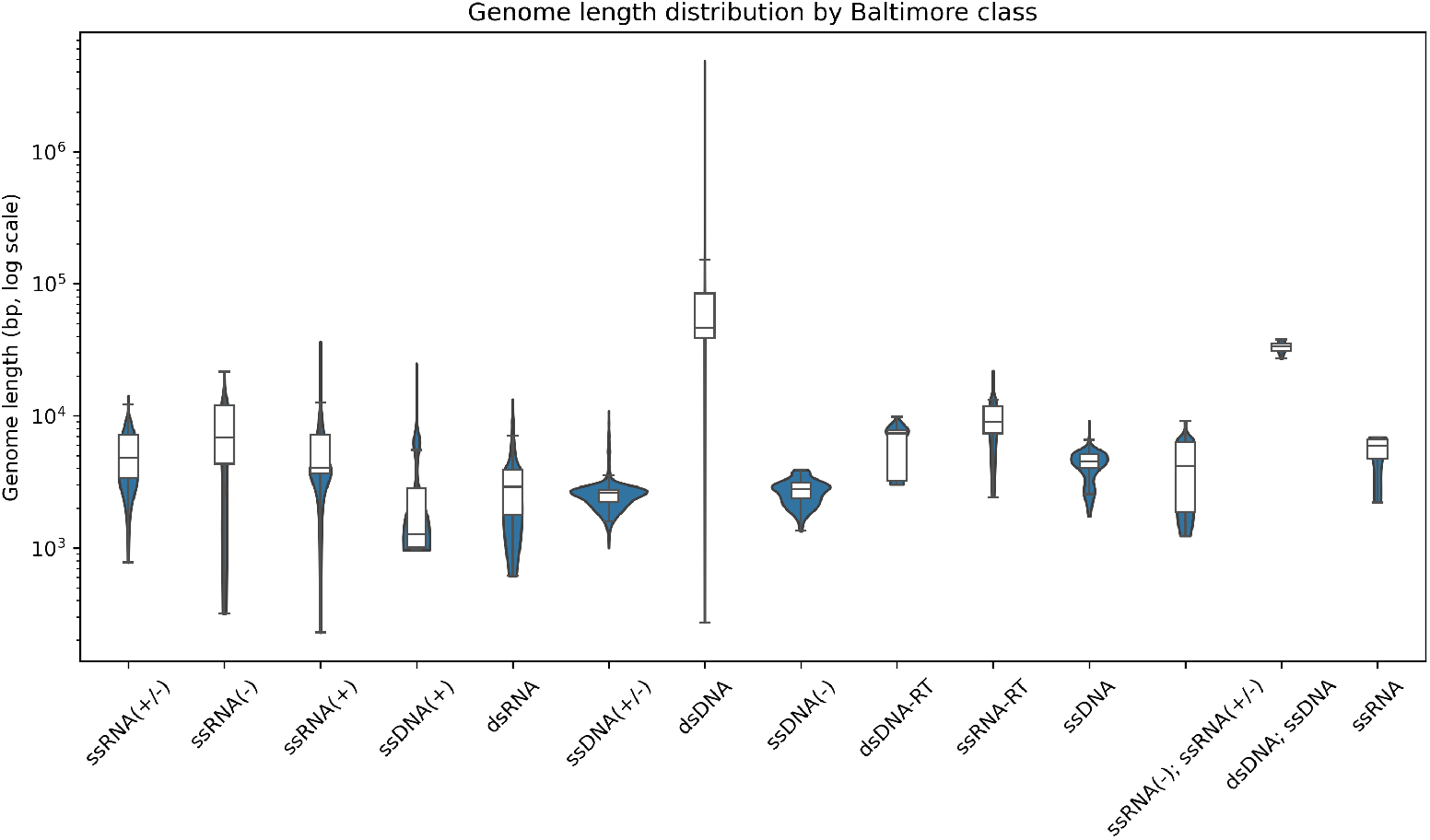
Distribution of viral genome lengths across Baltimore genome types. Violin plots represent the density of genome sizes (log scale), with overlaid boxplots indicating median and interquartile range.

#### S4.1 Genus-Level Performance Stratified by Genome Properties

Genus-level classification performance was further stratified by Baltimore genome type and by genome length. Detailed metrics for each Baltimore category are reported in Supplementary Table 7, while performance across genome length bins is summarised in Supplementary Table 8.

**Table 7.** Genus-level classification performance stratified by genome type.

| Genome | Accuracy | Balanced Acc. | Macro-F1 | N |
| --- | --- | --- | --- | --- |
| dsDNA | 0.941 | 0.893 | 0.879 | 7155 |
| dsDNA-RT | 0.914 | 0.646 | 0.505 | 151 |
| dsDNA;<br>ssDNA | 1.000 | 1.000 | 1.000 | 10 |
| dsRNA | 0.925 | 0.864 | 0.840 | 1116 |
| ssDNA | 0.831 | 0.579 | 0.530 | 255 |
| ssDNA(+) | 0.726 | 0.641 | 0.590 | 252 |
| ssDNA(+/-) | 0.769 | 0.461 | 0.405 | 2078 |
| ssDNA(-) | 0.410 | 0.173 | 0.114 | 166 |
| ssRNA | 1.000 | 1.000 | 1.000 | 8 |
| ssRNA(+) | 0.781 | 0.472 | 0.458 | 5534 |
| ssRNA(+/-) | 0.956 | 0.841 | 0.697 | 225 |
| ssRNA(-) | 0.824 | 0.592 | 0.564 | 2400 |
| ssRNA(-);<br>ssRNA(+/-) | 0.860 | 0.611 | 0.533 | 385 |
| ssRNA-RT | 0.926 | 0.733 | 0.614 | 68 |

**Table 8.** Genus-level classification performance stratified by genome length.

| Length Bin | Accuracy | Balanced Acc. | Macro-F1 | N |
| --- | --- | --- | --- | --- |
| 0–5kb | 0.745 | 0.401 | 0.375 | 8308 |
| 5–15kb | 0.879 | 0.754 | 0.729 | 4539 |
| 15–50kb | 0.970 | 0.926 | 0.902 | 3712 |
| 50–200kb | 0.963 | 0.900 | 0.887 | 2990 |
| ≥200kb | 0.909 | 0.839 | 0.789 | 254 |

## Footnotes

1 This analysis is post hoc on viruses with HMM hits; the score is not used during training or inference.

## Notes

### Competing Interest Statement

The authors have declared no competing interest.

## References

1. L. Call, S. Nayfach, and N. C. Kyrpides. Illuminating the virosphere through global metagenomics. Annual Review of Biomedical Data Science, 4(1):369–391, 2021.

2. A. P. Camargo, S. Roux, F. Schulz, M. Babinski, Y. Xu, B. Hu, P. S. Chain, S. Nayfach, and N. C. Kyrpides. Identification of mobile genetic elements with genomad. Nature biotechnology, 42(8):1303–1312, 2024.

3. A. C. Gregory, A. A. Zayed, N. Conceição-Neto, B. Temperton, B. Bolduc, A. Alberti, M. Ardyna, K. Arkhipova, M. Carmichael, C. Cruaud, et al. Marine dna viral macro-and microdiversity from pole to pole. Cell, 177(5):1109–1123, 2019.

4. Y. He, F. Zhou, J. Bai, Y. Gao, X. Huang, and Y. Wang. Vitax: adaptive hierarchical viral taxonomy classification with a taxonomy belief tree on a foundation model. Briefings in Bioinformatics, 26(1):bbaf041, 2025.

5. D. Hyatt, G.-L. Chen, P. F. LoCascio, M. L. Land, F. W. Larimer, and L. J. Hauser. Prodigal: prokaryotic gene recognition and translation initiation site identification. BMC bioinformatics, 11(1):119, 2010.

6. J.-Z. Jiang, W.-G. Yuan, J. Shang, Y.-H. Shi, L.-L. Yang, M. Liu, P. Zhu, T. Jin, Y. Sun, and L.-H. Yuan. Virus classification for viral genomic fragments using phagcn2. Briefings in bioinformatics, 24(1), 2023.

7. E. V. Koonin, V. V. Dolja, and M. Krupovic. The logic of virus evolution. Cell host & microbe, 30(7):917–929, 2022.

8. E. V. Koonin, M. Krupovic, and V. I. Agol. The baltimore classification of viruses 50 years later: how does it stand in the light of virus evolution? Microbiology and Molecular Biology Reviews, 85(3):10–1128, 2021.

9. E. V. Koonin, J. H. Kuhn, V. V. Dolja, and M. Krupovic. Megataxonomy and global ecology of the virosphere. The ISME journal, 18(1):wrad042, 2024.

10. T. Mihara, H. Koyano, P. Hingamp, N. Grimsley, S. Goto, and H. Ogata. Taxon richness of “megaviridae” exceeds those of bacteria and archaea in the ocean. Microbes and environments, 33(2):162–171, 2018.

11. D. Paez-Espino, I.-M. A. Chen, K. Palaniappan, A. Ratner, K. Chu, E. Szeto, M. Pillay, J. Huang, V. M. Markowitz, T. Nielsen, et al. Img/vr: a database of cultured and uncultured dna viruses and retroviruses. Nucleic acids research, 45(D1):gkw1030, 2016.

12. D. Paez-Espino, S. Roux, I.-M. A. Chen, K. Palaniappan, A. Ratner, K. Chu, M. Huntemann, T. B. K. Reddy, J. C. Pons, M. Llabrés, et al. Img/vr v. 2.0: an integrated data management and analysis system for cultivated and environmental viral genomes. Nucleic acids research, 47(D1):D678–D686, 2019.

13. J. C. Pons, D. Paez-Espino, G. Riera, N. Ivanova, N. C. Kyrpides, and M. Llabrés. Vpf-class: taxonomic assignment and host prediction of uncultivated viruses based on viral protein families. Bioinformatics, 37(13):1805–1813, 2021.

14. C. Riccardi, Y. Wang, S. Yooseph, and F. Sun. Bidirectional subsethood of shared marker profiles enables accurate virus classification. Microbiome, 13(1):170, 2025.

15. M. Unterer, M. Khan Mirzaei, and L. Deng. Gut phage database: phage mining in the cave of wonders. Signal Transduction and Targeted Therapy, 6(1):193, 2021.

